# A pesticide paradox: Fungicides indirectly increase fungal infections

**DOI:** 10.1101/156018

**Authors:** Jason R. Rohr, Jenise Browna, William A. Battaglin, Taegan A. McMahon, Rick A. Relyea

## Abstract

There are many examples where the use of chemicals have had profound unintended consequences, such as fertilizers reducing crop yields (paradox of enrichment) and insecticides increasing insect pests (by reducing natural biocontrol). Recently, the application of agrochemicals, such as agricultural disinfectants and fungicides, has been explored as an approach to curb the pathogenic fungus, *Batrachochytrium dendrobatidis* (*Bd*), which is associated with worldwide amphibian declines. However, the long-term, net effects of early-life exposure to these chemicals on amphibian disease risk have not been thoroughly investigated. Using a combination of laboratory experiments and analysis of data from the literature, we explored the effects of fungicide exposure on *Bd* infections in two frog species. Extremely low concentrations of the fungicides azoxystrobin, chlorothalonil, and mancozeb were directly toxic to *Bd* in culture. However, estimated environmental concentrations of the fungicides did not reduce *Bd* on Cuban tree frog (*Osteopilus septentrionalis*) tadpoles exposed simultaneously to any of these fungicides and *Bd*, and fungicide exposure actually increased *Bd*-induced mortality. Additionally, exposure to any of these fungicides as tadpoles resulted in higher *Bd* abundance and greater *Bd*-induced mortality when challenged with *Bd* post-metamorphosis, an average of 71 days after their last fungicide exposure. Analysis of data from the literature revealed that previous exposure to the fungicide itraconazole, which is commonly used to clear *Bd* infections, made the critically endangered booroolong frog (*Litoria booroolongensis*) more susceptible to *Bd*. Finally, a field survey revealed that *Bd* prevalence was positively associated with concentrations of fungicides in ponds. Although fungicides show promise for controlling *Bd*, these results suggest that, if fungicides do not completely eliminate *Bd* or if *Bd* re-colonizes, exposure to fungicides has the potential to do more harm than good. To ensure that fungicide applications have the intended consequence of curbing amphibian declines, researchers must identify which fungicides do not compromise the pathogen resistance mechanisms of amphibians.

## Introduction

Synthetic chemicals have had enormous value to society, such as by treating and curing diseases and revolutionizing agricultural production. However, they can also have unanticipated consequences (Pimentel et al. 1973). In fact, there are several examples where applications of chemicals to the environment have had exactly the opposite effects than were intended. For example, the application of fertilizers can destabilize crop herbivore dynamics resulting in larger herbivore outbreaks that, in some years, can result in zero crop yields, a phenomenon called the paradox of enrichment (Rosenzweig 1971). Similarly, the application of insecticides has, in some cases, actually increased pests by having more adverse effects on arthropod predators of the pests than on the pests themselves, thus adversely affecting natural biocontrol (Desneux et al. 2007, Douglas et al. 2015). Additionally, there are numerous examples where chemicals can have substantial non-target effects that can disrupt rather than enhance ecosystem functions and services (McMahon et al. 2012, Halstead et al. 2014, Staley et al. 2014). These examples highlight the need to comprehensively understand the complex effects that chemicals can have on ecosystems to avoid inadvertent and undesirable ramifications.

Recently, several researchers have explored the effects of pesticides on the chytrid fungus, *Batrachochytrium dendrobatidis* (*Bd*), a pathogen associated with worldwide amphibian declines (Kilpatrick et al. 2010). For example, studies have explored how *Bd* is affected by herbicides (Gahl et al. 2011, McMahon et al. 2013, Rohr et al. 2013, Buck et al. 2015, Jones et al. 2016), insecticides (Davidson et al. 2007, Gaietto et al. 2014, Buck et al. 2015, Jones et al. 2016), fungicides (Johnson et al. 2003, Woodhams et al. 2012, McMahon et al. 2013, Gaietto et al. 2014, Hudson et al. 2016), and agricultural disinfectants (Johnson et al. 2003, Bosch et al. 2015). Additionally, many researchers are exploring biocides as management tools to control *Bd* (Woodhams et al. 2011). For example, Hanlon et al. (Hanlon et al. 2012) showed that the common agricultural fungicide thiophanate-methyl cleared frogs of *Bd* and had positive effects on their health, and Woodhams et al. (Woodhams et al. 2011) highlighted several studies where fungicides and disinfectants were being explored as management tools. More specifically, Bosch et al. (2015) successfully eradicated *Bd* from a field site using a fungicide and an agricultural disinfectant, and Hudson et al. (2016) temporarily reduced fungal loads on amphibians in the wild using *in situ* exposure to a fungicide. Thus, chemicals, such as fungicides and disinfectants with fungicidal properties, show exciting promise for managing amphibian chytridiomycosis. However, the long-term, net effects of early-life exposure to these chemicals on amphibian disease risk have not been thoroughly investigated.

In addition to intentional exposure to products with fungicidal properties to manage *Bd*, amphibians are also regularly and unintentionally exposed to fungicides from direct overspray, drift, and run-off (Maltby et al. 2009, McMahon et al. 2011). Fungicides have experienced a greater increase in use in the last two decades than herbicides and insecticides. For example, in the US from 2004-2005, only 2% of all corn, soybean, and wheat fields were sprayed with fungicides, but this number increased to nearly 30% by 2009 (Belden et al. 2010). The soybean industry estimates that 60 to 70% of soybean seed planted in 2014 had a fungicide seed treatment, compared to 30% in 2008 and 8% in 1996 (http://unitedsoybean.org/article/six-things-farmers-should-know-about-seed-treatments/). Additionally, in the last 15 years in the US, an average of two new fungicides were registered for use each year, and in 2011 alone, the US Environmental Protection Agency reviewed the registration of 14 fungicides for 49 new uses, and all of the 14 fungicides are in use today (Battaglin et al. 2011).

Despite the widespread and increasing use of fungicides, there are few studies of the direct and indirect effects of fungicides on aquatic ecosystems (but see McMahon et al. 2011, McMahon et al. 2012) compared to the copious research on insecticides and herbicides (e.g. Relyea 2005, Rohr and Crumrine 2005, Rohr and McCoy 2010). For instance, a review of the indirect effects of pesticides on aquatic food webs found that only two out of the 150 research articles written from 1970-2002 addressed the use of a fungicide (Fleeger et al. 2003). Fungicides also are often very broad spectrum, affecting common and vital physiological processes, such as cellular respiration and immunity (Maltby et al. 2009). Additionally, modeling and monitoring suggest that fungicides might accumulate in freshwater habitats to levels above those considered safe for chronic exposure of some aquatic organisms (Deb et al. 2010).

The effects of pesticides on host-parasite interactions in particular can be extremely complex, having both positive and negative effects. As mentioned, several studies have shown that fungicides can be directly toxic to *Bd* (Hanlon and Parris 2012, McMahon et al. 2013). Chemical contaminants can also have negative effects on hosts by altering host immunity or parasite virulence (Relyea and Hoverman 2006, Rohr et al. 2006a, Jayawardena et al. 2016). Indeed, many pesticides are known to be immunomodulators (Voccia et al. 1999, Rohr et al. 2008b, Rohr and McCoy 2010, Rohr et al. 2015) and thus might have unfavorable effects on amphibian disease risk. Given the complexity of these direct and indirect effects, it is not surprising that studies exploring the effects of pesticides on *Bd* infections have produced mixed results (Davidson et al. 2007, Hanlon et al. 2012, McMahon et al. 2013, Buck et al. 2015, Hanlon et al. 2015, Jones et al. 2016).

Ultimately, what must be quantified is the sum of the positive and negative effects of contaminants to assess an overall or net effect (Rohr et al. 2008a). Additionally, net effects must be considered across life stages because exposure to chemicals early in life can have effects that persist into adulthood (e.g. Rohr and Palmer 2005, Rohr et al. 2006b). As an example, although the pesticide atrazine was directly toxic to *Bd* in culture (McMahon et al. 2013), the net effect of amphibian early-life exposure to this pesticide was an increase in *Bd*-induced mortality because it had adverse effects on frog tolerance of infections that occurred later in life (Rohr et al. 2013).

Here, we use a combination of experiments and analysis of data from the literature to explore the net effects of exposure to four common fungicides on *Bd* infections in two frog species. To test for direct effects, we exposed *Bd* in culture to three commonly used fungicides each at five ecologically relevant concentrations. To test for indirect and persistent effects of fungicides, we conducted a laboratory experiment where tadpoles were exposed to these three fungicides or a control and challenged with *Bd* during the fungicide exposure period and/or after metamorphosis. To compare our results on these three fungicides used commonly in agriculture to itraconazole, the fungicide commonly used to clear amphibians of *Bd* (Garner et al. 2009, Berger et al. 2010), we analyzed data from the literature on the effects of itraconazole on susceptibility to *Bd*. Finally, to evaluate the relevance of our results to natural settings, we tested whether fungicide levels in waterbodies and frog tissues were correlated with the prevalence of *Bd* infections.

We predicted that each fungicide would be directly toxic to *Bd*. Given that most of these fungicides are documented to suppress immune responses of vertebrates (Colosio et al. 1996, Corsini et al. 2006, McMahon et al. 2011), we also expected that exposure to each fungicide would alter amphibian-*Bd* interactions (indirect effect). The direction of the net effect of fungicide exposure, however, was challenging to predict *a priori*. Additionally, we predicted that if any fungicide delayed metamorphosis, it would increase *Bd*-loads on frogs by increasing the amount of time tadpoles were exposed to this predominantly aquatic fungus.

### Background on pathogen and fungicides

*Bd* is a pathogenic chytrid fungus that causes chytridiomycosis in many amphibians. It is a major contributor to global amphibian declines and has been found on six continents (Kilpatrick et al. 2010). *Bd* infects amphibians by colonizing keratin-containing body regions, infecting the mouthparts of anuran tadpoles and later spreading to the skin as the skin becomes keratinized during metamorphosis (McMahon and Rohr 2015). Infection does not usually cause tadpole mortality, but post-metamorphic anurans can be extremely susceptible to *Bd* (McMahon et al. 2013, Gervasi et al. 2017) because the infection disrupts osmotic and electrolyte balance that is controlled by their skin, which can eventually lead to cardiac arrest (Voyles et al. 2009). Cellular immunity is an important defense against *Bd* and despite *Bd* being immunosuppressive, amphibians can acquire immunological resistance to *Bd* that overcomes this immunosuppression (McMahon et al. 2014).

Four fungicides were studied here: azoxystrobin, chlorothalonil, mancozeb, and itraconazole. The first three fungicides rank among the top five in the United States based on usage (Grube et al. 2011). Azoxystrobin has experienced a recent increase in usage in the US in the last couple of decades to combat the emergence of soybean rust and is also used commonly on grain crops (Battaglin et al. 2011). Chlorothalonil is used on a wide variety of crop species, as well as residential and golf course turf (Caux et al. 1996), and is the fungicide most commonly used in Latin America, a place where *Bd* effects on amphibians have been severe (Ghose et al. 2014). Mancozeb is primarily applied to potatoes to reduce fungal pathogens and is also sprayed on other food crops (Maltby et al. 2009, Grube et al. 2011). Azoxystrobin and chlorothalonil inhibit cellular respiration, and mancozeb disrupts lipid metabolism (Caux et al. 1996, Maltby et al. 2009, Battaglin et al. 2011), and thus all three chemicals have modes of action that affect vital physiological processes of many organisms, from bacteria to vertebrate animals. The half-lives and estimated peak environmental concentrations (based on US EPA EXAMS-PRZM software) of azoxystrobin, chlorothalonil, and mancozeb are 11-17 d, 1 to 48 h, and 1-2 d and ~2 ppb, ~164 ppb, and ~58 ppb, respectively.

Itraconazole is not heavily used in agriculture but is used in the medical and veterinary fields and is probably the most commonly used fungicide to clear *Bd* infections of amphibians (KuKanich 2008, Garner et al. 2009, Berger et al. 2010). Its mode of action is to inhibit fungal-mediated synthesis of ergosterol, and it can also inhibit cytochrome P450, which is important in metabolizing potentially toxic compounds (KuKanich 2008).

## Materials and Methods

### Preparation of *Bd* inoculum

*Bd* inoculum was prepared by adding 1 mL of *Bd* stock (isolate SRS 812 isolated from a *Rana catesbeiana* in 2006 captured near the Savannah River Ecology Lab, SC, USA, passed through culture ~12 times) cultured in 1% tryptone broth, to a 1% tryptone agar plate. The plates were maintained at 23°C for approximately 1 week to allow *Bd* proliferation. Plates were inspected microscopically to verify that the zoospores were viable and then they were then flooded with ultrapure water to suspend the zoospores. Water from all the plates was homogenized to create the *Bd* positive (*Bd*+) solutions. Zoospore density was standardized among replicates and concentrations above the target concentration were diluted with ultrapure water. The *Bd* negative (*Bd-*) solution was created using the same method, except that no *Bd* was added to the 1% tryptone agar plates.

### Effects of fungicides on Bd in culture

To test for direct effects of the fungicides on *Bd* in culture, we used methods developed by McMahon et al. (2013). In a sterile, laminar flow hood, 10-mL glass test tubes were filled with *Bd*+ solution (total concentration: 3.8x10^4^ zoospores/mL), 1% tryptone broth, and one of four fungicide stocks (0.1x, 1.0x, 10x, and 100x EEC of each of the three fungicides) or two control treatments (water and acetone solvent controls). Each treatment (concentration of each fungicide) was replicated 5 times for a total of 25 experimental units per fungicide, plus 10 control replicates. Test tubes were maintained at 23°C for 10 d after which *Bd* quantities were quantified by counting a 10-µL aliquot of *Bd* with a hemocytometer.

### Effects of fungicides on Bd growth on frogs in the laboratory

The objectives of this experiment were to test for (1) the direct effects of fungicides on amphibians, (2) indirect effects of fungicides mediated by *Bd*, and (3) persistent effects of fungicide exposure on amphibian susceptibility to *Bd*. To accomplish these objectives, we collected Cuban tree frog tadpoles (*Osteopilus septentrionalis*) from a pond near the University of South Florida Botanical Gardens during the month of September 2012. We chose this frog species because it is common locally but is susceptible to chytrid infections (Rohr et al. 2013). In summary, we conducted a 4 x 2 x 2 study with exposure to one of four fungicide treatments during larval development (azoxystrobin, chlorothalonil, mancozeb, solvent control) crossed with *Bd* exposure or not during larval development crossed with *Bd* exposure or not after metamorphosis (see below; Table S1, Fig. S1).

Each individual tadpole was weighed and staged (Gosner 1960) before the start of the experiment; stages and weights were equally distributed across all treatments. Individual tadpoles were exposed to treatments in a 500-mL glass jar filled with 300 mL of artificial spring water (ASW, Cohen et al. 1980). Throughout the experiment, animals were fed fish flakes and Sera Micron *ad libitum*.

The four fungicide treatments were an acetone control or the EEC of azoxystrobin (2.06 µg/L), chlorothalonil (164 µg/L, we used the concentration of 30.0 µg/L), or mancozeb (57.6 µg/L). The treatments were applied to the water in each jar at the start of the experiment. These treatments were re-applied with every water change, which occurred weekly until metamorphosis or up to 12 weeks.

The *Bd* exposures occurred during the tadpole stage (simultaneous exposure with the fungicide treatments), after metamorphosis (delayed exposure; after fungicide treatments), both during the tadpole stage and after metamorphosis (double exposure; during and after fungicide treatments), or not at all (sham-exposed during both life stages). For the tadpole exposures, a 1-mL *Bd* inoculum (2.88x10^6^ - 4.25x10^6^ zoospores/mL) was added to the appropriate jar during weeks 1 and 3 of the experiment. All animals that were randomly assigned not to receive *Bd* during a given exposure period received a sham *Bd* exposure (see *Preparation of Bd stocks*).

The tadpoles were checked daily for metamorphosis or mortality. All animals that metamorphosed were swabbed (snout to vent and down each leg 5 times each), weighed, and maintained individually in 1-L plastic deli cups at 23°C. Each post-metamorphic juvenile frog received vitamin- and mineral-dusted crickets *ad libitum* and weekly changes of wet papers towel. All animals that died were swabbed, weighed, and preserved. After 12 weeks, any tadpoles that had not metamorphosed were swabbed (mouthparts only; see McMahon and Rohr 2015) and euthanized using 0.5% MS-222. Animals were then weighed, staged, and snout-vent length (SVL) was measured. The mouthparts were removed and stored in 95% ethanol to later measure *Bd* loads.

At week 22, all metamorphic frogs were exposed to either 1mL of *Bd+* inoculum (1.69x10^6^ zoospores/mL) or 1mL of the *Bd*- inoculum. For animals that were exposed only to a fungicide or control treatment as tadpoles, this was their first exposure to *Bd*; for the simultaneous *Bd*-fungicide treatment animals, this was the second *Bd* exposure. Five weeks after post-metamorphic exposure to *Bd* or the sham inoculum, the right hind limb (15 times hip to toe) of all frogs were swabbed. To optimize *Bd* growth, all frogs were moved to 17°C at week 32; all frogs were swabbed (15 times hip to toe) and then euthanized during week 38 (Fig. 2b).

Both swabs and tissues from the mouthparts of tadpoles provided DNA to quantify *Bd* abundance. *Bd* DNA was amplified through qPCR to calculate *Bd* infection abundance. All qPCR was performed according to Hyatt et al. (2007), using StepOne™ Real-Time PCR System (qPCRStepOne™).

We initially had 10 replicate frogs for each of these 16 treatments (160 frogs total). However, if a frog was assigned to receive a *Bd* exposure after metamorphosis and it died before this exposure, the animal was re-assigned to the appropriate treatment. For example, if a frog was assigned to receive *Bd* exposure both before and after metamorphosis but died before receiving the second *Bd* exposure after metamorphosis, it was shifted from the double *Bd* exposure treatment to the *Bd* before metamorphosis-only treatment. Likewise, if a frog was assigned to only receive *Bd* after metamorphosis and it died before receiving this treatment, it was reassigned to the no *Bd* before or after metamorphosis treatments. Hence, this had the effect of increasing sample sizes in some groups while decreasing the sample size in others. Sample sizes ended up as follows: simultaneous exposure *n*=53; delayed exposure *n* = 35; double exposure *n*=25; sham-exposure *n*=42.

### Effects of the fungicide itraconazole on Bd growth on frogs in the laboratory

Itraconazole is probably the most commonly used fungicide to clear frogs of *Bd* because of the existence of established amphibian application protocols (Garner et al. 2009, Berger et al. 2010) and thus is extremely relevant to the amphibian-*Bd* system. To evaluate whether itraconazole had similar effects as azoxystrobin, chlorothalonil, and mancozeb and to provide a test of the effects of fungicides on *Bd* infections by a completely independent laboratory (adding to the weight of evidence), we searched the literature for studies that exposed amphibians to itraconazole and then challenged them with *Bd*. Cashins et al. (2013) conducted a study on the critically endangered booroolong frogs (*Litoria booroolongensis*) that satisfied this search criteria. In this study, Cashins et al. had four groups of frogs. On day one of the experiment, Group 1 was exposed to *Bd* and groups 2-4 were not. On day 30, Groups 1 and 2 were exposed to itraconazole to clear the initial *Bd* infection and to control for the itraconazole exposure, respectively. On day 110, Groups 1-3 were exposed to *Bd* and Group 4 received a sham control. On day 179, *Bd* prevalence was quantified. Sample sizes are provided in Cashins et al. (2013) but were >10 independent frogs per treatment group. In summary, Group 1 was exposed to itraconazole and *Bd* twice, Group 2 was exposed to itraconazole and *Bd* once, Group 3 was never exposed to itraconazole but was exposed to *Bd* once, and Group 4 was not exposed to either itraconazole or *Bd*. Cashins et al. (2013) focused on how previous exposure to *Bd* affected host resistance upon a second exposure but did not test for the effects of itraconazole on *Bd* prevalence in their study. This statistical test is described in the *Data analysis* section below.

### Effects of the fungicides on Bd prevalence in the field

To investigate for an association between fungicide exposure and *Bd* infections, we collected water, bed sediment, *Bd*, and frog tissue samples concurrently from 21 sites in seven states (CA, CO, GA, ID, LA, ME, and OR) in 2009 and 2010 and quantified *Bd* and fungicide concentrations. Details on the site locations, characteristics, and sampling design are provided in detail in Battaglin et al. (2016) and Smalling et al. (2015) and thus and thus are only briefly covered here. Approximately 1 L of water and a stainless steel scoop of bed sediment were collected from each site to quantify pesticide or pesticide degradates. One to 15 adult frogs were collected at each site (see Results for details on frogs species collected) by hand or net and swabbed for *Bd* (Hyatt et al. 2007). The frogs were euthanized, wrapped in aluminum foil, and then placed in a freezer for later whole-body analysis of pesticides and pesticide degradates (Battaglin et al. 2016).

Filtered water samples were analyzed for a suite of pesticides and pesticide degradates by extracting onto a solid phase extraction cartridge, spiking the samples with a recovery surrogate, eluting the cartridge with ethyl acetate, adding a deuterated internal standard, and analyzing the extracts on an Agilent (Santa Clara, CA) 7890 gas chromatograph coupled to an Agilent 5975 (Folsom, CA) mass spectrometer. Data for all pesticides were collected in selective ion monitoring mode with each compound having one quantifier ion and one to two qualifier ions. Wet sediments (10 g) were analyzed similarly to the water samples with the following exceptions. The sediment was homogenized with sodium sulfate, extracted using pressurized liquid extraction, dried over sodium sulfate, reduced, and sulfur was removed by gel permeation chromatography. Samples were subjected to a clean-up method with 6% deactivated Florisil. Frog tissue samples were analyzed similarly to the water samples with the following exceptions. Individual whole frogs (3–5 g) were thawed, homogenized with sodium sulfate (Na_2_SO_4_) using a mortar and pestle, extracted three times with dichloromethane using pressurized liquid extraction, dried over Na_2_SO_4_, and reduced to 1 mL. Ten percent by volume of each raw extract was allowed to evaporate to a constant weight in a fume hood for gravimetric lipid determination to the nearest 0.001 g using a microbalance. A majority of the lipid was removed using gel permeation chromatography followed by 6% deactivated Florisil previously activated at 550 °C for 16 h (Battaglin et al. 2016).

### Data analysis

All statistical analyses were conducted using R statistical software. For the study that examined *Bd* responses to three fungicides, we conducted analyses on each of the three fungicides separately. We tested for a relationship between fungicide concentration and *Bd* abundance (rounded zoospore equivalents) using a negative binomial distribution (function: glmmadmb, package: glmmadmb). We tested for differences among the concentrations using a sequential Bonferroni adjustment.

For the laboratory study on frogs, we used a generalized linear model with a binomial error distribution to determine whether the fungicide treatments, *Bd* treatments, and their interaction significantly affected whether a frog lived or died during the experiment, and used a factorial ANOVA to evaluate how treatments affected log mass at and time to metamorphosis. Additionally, we conducted two analyses on *Bd* abundance. First, we compared the loads of frogs exposed to *Bd* for the first time across fungicide treatments to see if simultaneous exposure to *Bd* and fungicides resulted in different fungal loads than early-life exposure to fungicides and later-life exposure to *Bd*. For frogs exposed simultaneously to *Bd* and fungicide treatments, the response variable was *Bd* load at metamorphosis or *Bd* load at death if they did not reach metamorphosis. For frogs exposed to *Bd* after they were exposed to fungicide treatments (i.e. after metamorphosis), we used the mid-survey (experimental day 155) *Bd* load or *Bd* load at death if they did not reach the mid-survey swabbing. Second, we compared the *Bd* load of post-metamorphic frogs receiving their first and second exposures to *Bd* using the mid-survey swabs. In all analyses treating *Bd* abundance on the frogs as a response, we included fungicide treatments, timing of *Bd* exposures, their interaction, and initial mass (unless it was not significant) as fixed effects. We analyzed *Bd* abundance using a zero-inflated negative binomial model (it was a better fit [AIC=1009] than the negative binomial model [AIC=1038]; function: zeroinfl, package: pscl).

To test whether the fungicide itraconazole was immunosuppressive to booroolong frogs in the Cashins et al. (2013) study, we applied a Chi square analysis to their data to compare *Bd* prevalence of frogs previously exposed to itraconazole or not.

Finally, to evaluate how fungicides affected *Bd* infections in the field, we used multiple regression with a binomial error distribution (function: glm, package stats) to test how the concentration of fungicides in frog tissues, sediment, and water affected *Bd* prevalence. We did not analyze *Bd* abundance data because we did not know the time course of infection for each frog, and thus we conservatively focused on prevalence. However, we did evaluate whether any differences among frog species in their fungicides or *Bd* loads could account for any detected patterns between fungicides and *Bd* prevalence. All *p*-values were calculated using log-likelihood ratio tests (function: Anova, package: car).

## Results

### Effects of fungicides on *Bd* in culture

Relative to the acetone control, all tested concentrations of azoxystrobin and chlorothalonil (Concentration: *X^2^*=19.96, *p<*0.001 and *X^2^*=15.22, *p<*0.001, respectively; Table 1) and all tested concentrations of mancozeb except the lowest concentration (Concentration: *X^2^*=6.81, *p=*0.009; Table 1) significantly reduced *Bd* abundance in culture relative to the controls.

**Table 1.** Effects of various concentrations of the fungicides azoxystrobin, chlorothalonil, and mancozeb on the abundance on *Batrachochytrium dendrobatidis* zoospores cultured in welled plates.

| Concentration (µg/L) | Mean log<sub>10</sub> zoospores/mL<sup>a</sup> | Standard error | Sequential Bonferroni<sup>b</sup> |
| --- | --- | --- | --- |
| Azoxystrobin |  |  |  |
| 0.000 | 5.65 | 0.15 | a |
| 0.002 | 1.03 | 1.03 | b |
| 0.020 | 0.00 | 0.00 | b |
| 0.206 | 2.27 | 0.94 | b |
| 2.060 | 0.00 | 0.00 | b |
| 20.600 | 0.00 | 0.00 | b |
| Chlorothalonil |  |  |  |
| 0.000 | 5.65 | 0.15 | a |
| 0.030 | 0.85 | 0.85 | b |
| 0.300 | 1.04 | 1.04 | b |
| 3.000 | 1.53 | 0.95 | b |
| 30.000 | 1.75 | 1.08 | b |
| 300.000 | 0.00 | 0.00 | b |
| Mancozeb |  |  |  |
| 0.000 | 5.65 | 0.15 | a |
| 0.057 | 4.81 | 0.45 | a,b |
| 0.570 | 1.04 | 1.04 | b |
| 5.700 | 3.70 | 0.99 | b |
| 57.600 | 0.00 | 0.00 | c |
| 576.000 | 0.68 | 0.68 | c |

<sup>a</sup>These means are from five independent samples per concentration.
<sup>b</sup>Concentrations that do not share a letter are significantly different from one another based on post hoc tests among concentrations within a fungicide.

### Effects of fungicides on Bd growth on frogs in the laboratory

There were no significant effects of the fungicides, *Bd* treatments, or their interactions on time to metamorphosis or mass at metamorphosis (Tables S1 & S2). For the *Bd* analyses, we first tested for an effect of fungicide treatments on *Bd* abundance on frogs that were exposed to *Bd* for the first time, which included frogs exposed to *Bd* and the fungicide treatments simultaneously and frogs exposed to fungicide treatments as tadpoles but to *Bd* an average of 71 days after metamorphosis. In these analyses, frogs were not all swabbed at the same time because they metamorphosed or died at different times. Duration of time that *Bd* grew on the frogs was never a significant predictor in the models (*X^2^*=2.97, *p*>0.225). However, whether a frog was swabbed at death was always a significant predictor (*p*<0.05) because frogs that died from infection had, on average, more *Bd* than frogs swabbed while alive. Therefore, we included whether a frog was swabbed at death as a categorical factor rather than time for *Bd* growth in each model. Initial mass was included in most models because it was generally a negative predictor of *Bd* intensity.

In these analyses, mean *Bd* abundance on frogs never differed significantly among the three fungicide treatments, regardless of whether *Bd* exposure occurred simultaneous with the fungicide exposure (*X^2^*=2.65, *p*=0.618) or later in life (*X^2^*=0.61, *p*=0.961; Table S3). In addition, exposure to any of the three fungicides resulted in significantly more *Bd* on frogs compared to frogs exposed to the solvent control (Fungicide main effect: *p*<0.05; Table S3). Hence, we pooled the fungicides together for ease of interpretation and to increase statistical power. There was no significant effect of the fungicides on *Bd* abundance when the fungicide and *Bd* exposures occurred simultaneously as tadpoles (Fig. 1A); however, when the *Bd* exposure occurred after metamorphosis, an average of 71 days since the previous exposure to fungicides, the previous fungicide exposure caused a nearly 3.5-fold increase *Bd* abundance on frogs relative to the solvent controls (Fungicide x timing of *Bd* exposure: *X^2^*=25.98, *p*<0.001; Fig. 1A).

**Fig. 1.**
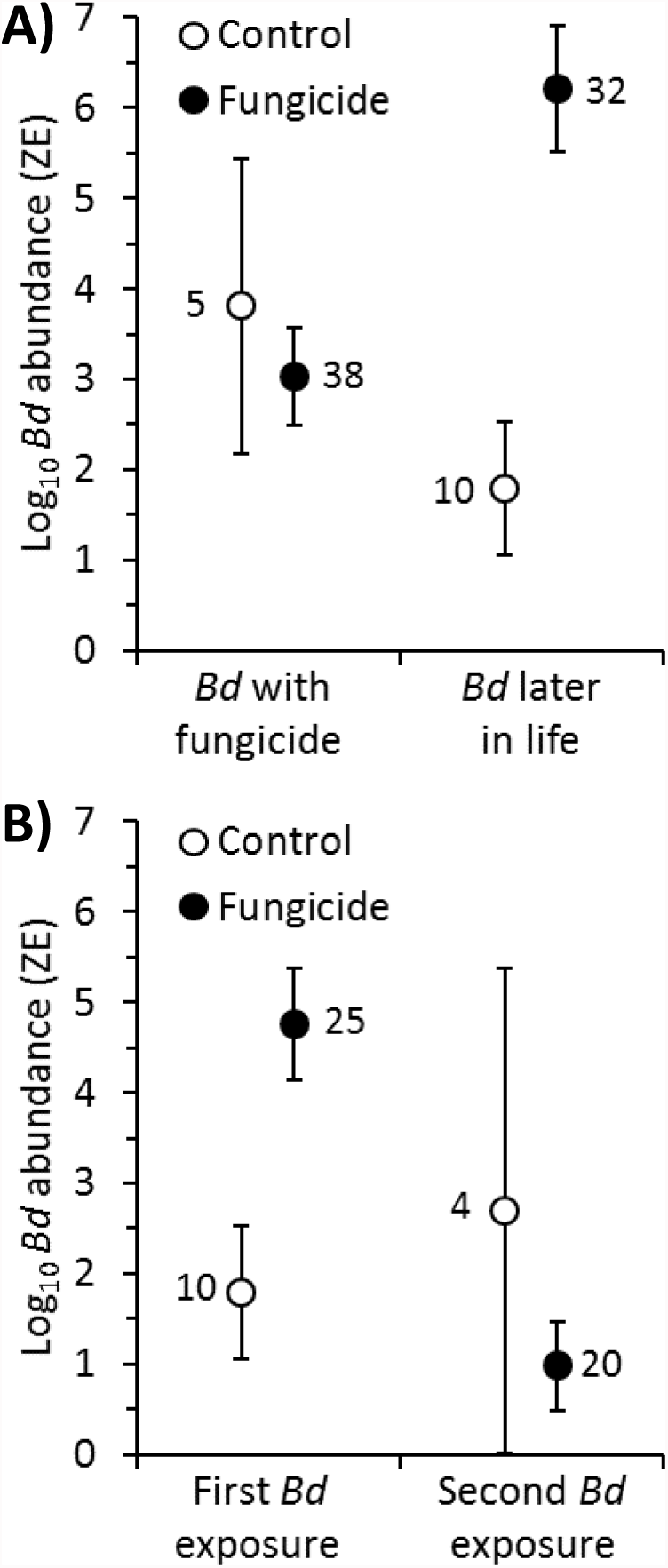
Effects of fungicide treatments on mean (± 1 SE) log10 *Batrachochytrium dendrobatidis* (*Bd*) abundance on Cuban tree frogs **A)** exposed to *Bd* for the first time as tadpoles simultaneous with the fungicide treatments versus for the first time as post-metamorphic frogs, 71 days after the previous exposure to fungicide treatments (Fungicide x timing of *Bd* exposure: *X^2^*=25.98, *p*<0.001), and **B)** exposed to *Bd* for the first versus second time as a post-metamorphic frog (Fungicide x number of *Bd* exposures: *X^2^*=15.47, *p*<0.001). There were no statistically significant differences among the three tested fungicides (azoxystrobin, chlorothalonil, mancozeb) and thus they were pooled for subsequent analyses to increase statistical power and facilitate visualization. Numbers next to data points are associated sample sizes (which vary because of mortality and re-assignment of frogs to appropriate treatments if they died before receiving their assigned *Bd* exposure; see text for details).

We also compared the *Bd* load of post-metamorphic frogs receiving their first and second exposures to *Bd*. These analyses revealed a significant two-way interaction between the fungicide treatment and number of *Bd* exposures (*X^2^*=15.47, *p*<0.001). Frogs exposed to fungicide and *Bd* for the first time had higher *Bd* loads than frogs exposed to solvent and *Bd* for the first time (*X^2^*=12.85, *p*=0.002; Fig. 1B, Table S4). However, the opposite trend occurred on the second exposure. Frogs exposed to fungicide and *Bd* for the second time had lower *Bd* loads than frogs exposed to the solvent and *Bd* for the second time, although not significantly so because of low sample size and high variability in the control (*X^2^*=2.95, *p*=0.229; Fig. 1B, Table S4).

For the analyses on survival, we conducted binomial survival analyses that assessed how treatments affected the probabilities of surviving the length of the experiment. In these analyses, mean mortality of frogs exposed to the three fungicides never differed, regardless of whether *Bd* exposure occurred simultaneous with the fungicide exposure (*X^2^*=1.33, *p*=0.515), later in life (delayed) (*X^2^*=0.23, *p*=0.892), or both (double exposure; *X^2^*=2.35, *p*=0.309). However, for the simultaneous and delayed exposure treatments, each fungicide caused significantly greater mortality than the solvent control (*p*<0.04). Hence, we again pooled the fungicides for ease of interpretation and to increase statistical power.

We compared *Bd*-induced mortality in the fungicide versus control treatments for frogs exposed to *Bd* for the first time. Despite not having significantly more *Bd* than controls, tadpoles exposed simultaneously to fungicide and *Bd* had greater mortality than tadpoles exposed to solvent, fungicides alone, and *Bd* alone, resulting in a significant interaction between fungicide and *Bd* treatments (Fungicide\**Bd*: *X^2^*=6.47, *p*=0.011; Fig. 2A). A similar but more pronounced pattern was observed when the *Bd* exposure occurred after metamorphosis, there was significantly greater *Bd*-induced mortality if frogs were previously exposed to fungicide than solvent control (Fungicide\**Bd*: *X^2^*=6.28, *p*=0.012; Fig. 2A).

**Fig. 2.**
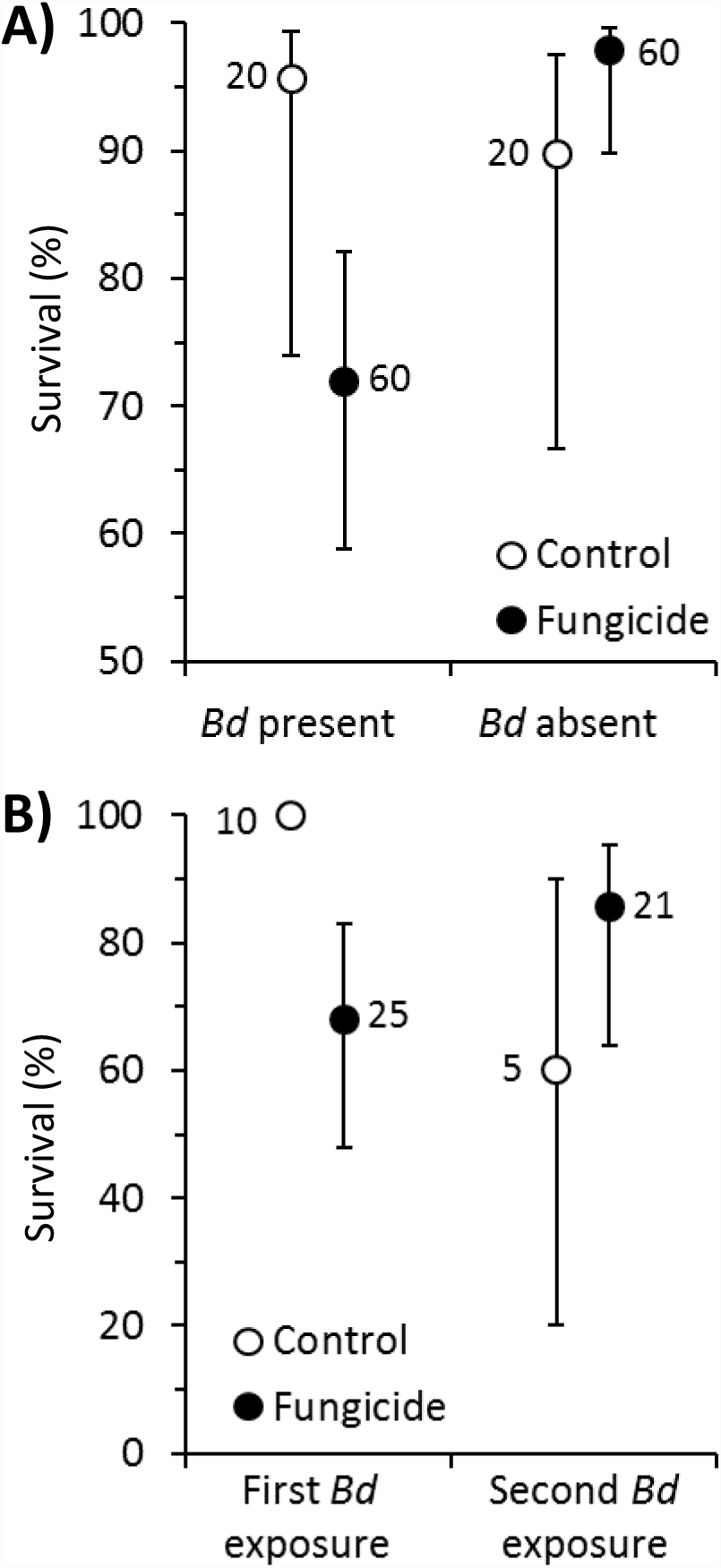
Mean (± 95% CI) survival of Cuban tree frogs exposed **A)** to fungicide and *Batrachochytrium dendrobatidis* (*Bd*) treatments simultaneously as tadpoles (Fungicide x *Bd*: *X^2^* = 6.47, *P*=0.011) or **B)** to fungicides treatments as tadpoles and then to *Bd* for the first or second time as a post-metamorphic frog (a mean of 71 days after their last exposure to fungicide; Fungicide x number of *Bd* exposures: *X^2^* = 6.89, *P*=0.009). There were no statistically significant differences among the three tested fungicides (azoxystrobin, chlorothalonil, mancozeb) and thus they were pooled for subsequent analyses to increase statistical power and facilitate visualization. Numbers next to data points are associated sample sizes (which vary because of mortality and re-assignment of frogs to appropriate treatments if they died before receiving their assigned *Bd* exposure; see text for details).

We also compared the *Bd*-induced mortality of post-metamorphic frogs receiving their first and second exposures to *Bd*. If metamorphs were exposed to *Bd* for the first time, there was greater *Bd*-induced mortality among frogs previously exposed to fungicide than solvent control (*X^2^*=6.28, *p*=0.012, Fig. 2B). However, similar to our *Bd* abundance results, the opposite trend was observed when metamorphs were exposed to *Bd* for the second time; there was greater *Bd*-induced mortality among frogs previously exposed to solvent control than fungicides, although not significantly so (*X^2^*=1.51, *p*=0.220; Fig. 2B). This resulted in a significant interaction between fungicide treatment and number of *Bd* exposures (*X^2^*=6.90, *p*=0.009) because there was a significant effect during the first exposure and not during the second exposure.

### Effects of the fungicide itraconazole on Bd growth on frogs in the laboratory

Booroolong frogs previously exposed to itraconazole in the Cashins et al. (2013) study had significantly higher *Bd* prevalence when exposed to *Bd* later in life (91%, 10/11 frogs) than frogs not previously exposed to itraconazole (50%, 14/28 frogs; *X^2^*=5.58, *p*=0.018).

### Effects of the fungicides on Bd growth on frogs in the field

Twelve different fungicides were detected in the 21 wetlands that were sampled (Table S5). The three most common fungicides detected in sediment were pyraclostrobin (52%), chlorothalonil (33%), and tebuconazole (24%), and the only fungicide detected from water samples at multiple sites (19%) was azoxystrobin (Table S5). Given the infrequency in which individual fungicides were detected, we combined fungicide concentrations for the analyses.

A total of 138 frogs were swabbed for *Bd* and also had fungicides quantified from their tissues. Of these frogs, 54 were positive for *Bd* and 72 had quantifiable levels of fungicides in their tissues. Only two wetlands had detectable levels of fungicides in the water column, whereas all but three had detectable levels in sediment (Table S5). Given how few wetlands had detectable fungicide in the water column, we chose to simply add the water column and sediment concentrations to reflect overall environmental exposure, but admit that much of this exposure appears to be through sediment. *Bd* prevalence on frogs was not significantly related to the concentration of fungicides in frog tissues (*X^2^*=0.03, *p*=0.858, Nagelkerke Index=0.001, Fig. 3A), but was positively related to the concentration of fungicides (sediment plus water) in waterbodies (*X^2^*=7.10, *p*=0.008, Nagelkerke Index=0.069, Fig. 3B). Two genera of frogs were captured, *Pseudacris* (65%) and *Rana* (35%; Fig. S2). Five species were captured but 65% were just two species, *Pseudacris regilla* and *P. triseriata* (Fig. S2). Loads of *Bd* were higher for genus *Pseudacris* than *Rana*, but *Pseudacris* was less likely to be found at sites with high concentrations of fungicides (Fig. S2). Thus, differences in *Bd* loads across genera do not seem capable of accounting for the detected positive relationship between fungicides and *Bd* (Fig. 3).

**Fig. 3.**
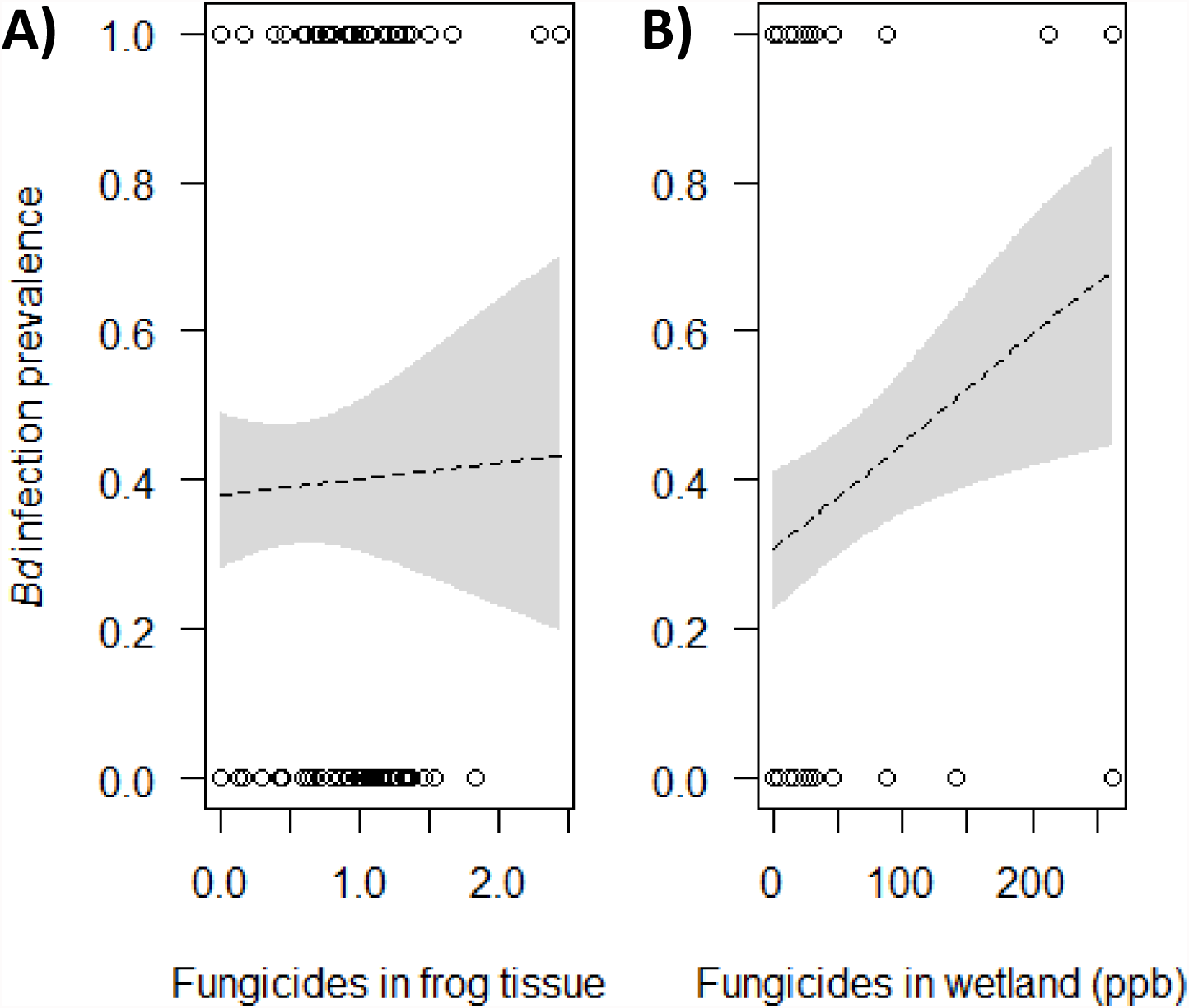
Relationship between the prevalence of *Batrachochytrium dendrobatidis* (*Bd*) infections of 138 frogs and the concentration of fungicides in frog tissues (log10(ppb+1)- transformed, *X^2^*=0.03, *p*=0.858, **A**) and in wetlands (sediment plus water, *X^2^*=7.10, *p*=0.008, **B**). Shown are logistic regression plots and associated 95% confidence bands (shaded).

## Discussion

When tadpoles were exposed to both a fungicide and *Bd* simultaneously in the laboratory, none of the three fungicides affected *Bd* loads on the frogs compared to the no-fungicide control. In contrast, frogs exposed to *Bd* as metamorphs, an average of 71 days since any fungicide exposure, had significantly greater *Bd* abundance and greater *Bd*-induced mortality than frogs exposed to the solvent control. The type of applied fungicide did not matter with all samples exhibiting the same reaction to the three fungicides, azoxystrobin, chlorothalonil, and mancozeb. Research from a completely independent laboratory on the critically endangered booroolong frog found exactly the same results for itraconazole (Cashins et al. 2013), the most commonly used fungicide to clear frogs of *Bd* (Garner et al. 2009, Berger et al. 2010). Booroolong frogs previously exposed to itraconazole had significantly greater *Bd* prevalence when challenged with *Bd* later in life than frogs not previously exposed to itraconazole. Hence, despite four commonly used fungicides being directly toxic to *Bd* (see Table 1), all paradoxically increased *Bd* infections by having persistent adverse effects on frog resistance to this fungal pathogen.

Interestingly, the increase in *Bd* abundance and *Bd*-induced mortality associated with fungicide exposure in our laboratory study was greater if we exposed the frogs to *Bd* an average of 71 days after the fungicide exposure than if we exposed the frogs to *Bd* and fungicide simultaneously. This result is probably a product of two factors. First, the culture experiment demonstrated that all three tested fungicides are directly toxic to *Bd* and thus almost certainly reduced the abundance of *Bd* on frogs when the two occurred simultaneously. However, any direct toxicity to *Bd* was clearly not as strong as the adverse effect of the fungicides on the fungal defenses of the frogs. Second, *Bd* is believed to consume keratin, which is only found on the mouthparts of tadpoles but is throughout the skin of metamorphs (McMahon and Rohr 2015). Hence, *Bd* might also be able to proliferate more rapidly after than before metamorphosis, amplifying the fungicide effect. Given that the fungicides did not affect timing of or size at metamorphosis, these traits seem unlikely to explain any observed effects.

The observed persistent effects of early-life exposure to fungicides on infectious disease risk are consistent with several previous toxicological studies. Several pesticides have been shown to cause changes in host-parasite dynamics (Relyea and Hoverman 2006, Rohr et al. 2006a, Rohr and McCoy 2010) and have delayed effects on host behavior, growth, physiology, and survival (e.g. Rohr and Palmer 2005, Rohr et al. 2006b, Jones et al. 2009, Rohr et al. 2013). Similar to our findings, other studies have shown that fungicides can be directly toxic to *Bd* (Hanlon and Parris 2012, McMahon et al. 2013). In contrast to our work, some of these previous studies revealed that fungicide exposures can actually reduce *Bd* growth rates on frogs, but these previous studies did not test for the effects of sequential fungicide and *Bd* exposures and did not test for persistent adverse effects of early-life exposure to fungicides (Hanlon et al. 2012, Hanlon and Parris 2012, McMahon et al. 2013, Hanlon et al. 2015).

There are several potential mechanisms by which early-life exposure to chemicals can have persistent effects on infectious disease risk. Pesticide exposure early in life can induce stress responses, elevating cortisol, corticosterone, or other stress-related hormones; chronic levels of these hormones have been associated with persistent immunomodulation (Martin et al. 2010, McMahon et al. 2011, McMahon et al. 2017). Chemical contaminants have also been shown to disrupt the microbiome of hosts. The gut microbiome has been linked to immune development in vertebrates (Hooper et al. 2012) and the skin microbiome in amphibians has been shown to inhibit the growth of *Bd* (Bletz et al. 2013). Two recent studies revealed that tadpole exposure to chemical contaminants reduced their gut and skin microbiota and reductions in gut microbiota were associated with reduced resistance to skin-penetrating gut nematodes and *Bd* later in life (Knutie et al. in review, Knutie et al. in revisions). Understanding the mechanisms by which pesticides cause long-term impacts on host defenses will be necessary to improve the design of pesticides.

Interestingly, the pattern of higher *Bd* abundance in fungicide- than solvent-exposed frogs was not apparent upon a second exposure to *Bd*. Frogs exposed to fungicides and to *Bd* for the second time had similar *Bd* loads as control frogs and had lower *Bd* loads than fungicide-exposed frogs exposed to *Bd* for the first time when exposed at the same life stage (Fig. 1B). This pattern was most likely caused by *Bd*-induced mortality. Frogs exposed to fungicide and *Bd* for the first time had significantly higher mortality than frogs exposed to solvent and *Bd* for the first time. In fact, *Bd* only caused significant mortality when frogs were exposed to fungicides. Hence, the most *Bd*-susceptible individuals were not available to be exposed to *Bd* for a second time in the fungicide treatments but were available for exposure to *Bd* a second time in the control treatment. Consequently, selection is a likely explanation for the change in *Bd* abundance and mortality patterns across fungicide treatments between the first and second *Bd* exposures (Rohr et al. 2008a).

Despite the broad spectrum nature of many fungicides (Maltby et al. 2009), we did not detect strong direct effects of the tested fungicides on Cuban tree frogs in our laboratory experiments. The fungicides alone did not affect survival or timing of or size at metamorphosis. Rather, most of the adverse or beneficial effects of fungicides were only apparent in the presence of *Bd.* The fungicides reduced *Bd* growth on frogs when the exposures occurred simultaneously. However, they tended to increase *Bd-*induced mortality regardless of the timing of exposures. These findings emphasize the importance of considering the effects of contaminants within a community context (Relyea and Hoverman 2006, Rohr et al. 2006a), quantifying the net effects of contaminants (the sum of the beneficial and adverse effects) (Rohr et al. 2008a), and testing for delayed or persistent effects of chemicals (Rohr and Palmer 2005, Rohr et al. 2006b, Jones et al. 2009, Rohr and Palmer 2013).

The patterns we observed in nature were consistent with our laboratory findings because fungicides were generally associated with greater rather than less *Bd*. In our field survey, we revealed that the greater the concentration of fungicides in a given wetland, the greater the prevalence of chytrid fungal infections in frogs (Fig. 3B). In contrast, fungicide concentrations at the level of individual frogs were less predictive of *Bd* prevalence (Fig. 3A), perhaps because of individual-level variation in susceptibility, exposure, and timing and duration of infections, variation that is reduced at the level of the wetland. In addition to reflecting current exposure, detectable levels of fungicides in wetlands must also reflect some level of previous fungicide exposure, which, according to our laboratory experiments, can persistently compromise host resistance to *Bd*. Importantly, given that *Bd* can be quite persistent in the presence of ample hosts and that fungicides often degrade rapidly in the environment, exposure to fungicide followed by exposure to *Bd* is almost certainly more frequent than simultaneous exposure to the two factors, suggesting that persistent adverse effects of previous fungicide exposure might be common. Additionally, our field patterns revealed that fungicides were rarely detected in the water column but were regularly detected in sediments, suggesting that adsorption might be important for many fungicides and that benthos-dwelling tadpole species might have higher fungicide exposure than species found more commonly in other microhabitats. Additional field data and field manipulations would be invaluable in determining the absolute magnitude of these effects and the specific fungicides that are driving these patterns.

Recently, there have been some exciting findings that suggest applications of fungicides or disinfectants with fungicidal properties might be an effective tool for curbing amphibian declines associated with *Bd* (Bosch et al. 2015, Hudson et al. 2016). Bosch et al. (2015) used fungicides and agricultural disinfectants to successfully eradicate *Bd* from a field site and Hudson et al. (2016) temporarily reduced fungal loads on amphibians in the wild using *in situ* exposure to fungicides. However, our results show that many fungicides can also have adverse effects, such as persistently compromising amphibian defenses against pathogens. Thus, fungicides might work well if they completely eliminate *Bd* from the environment and if *Bd* is unlikely to return soon after. However, if a fungicide application does not eradicate *Bd*, if it does eradicate *Bd* but *Bd* re-colonizes, or if fungicide-induced suppression of host defenses affects resistance to other virulent pathogens in the environment (bacteria, viruses, macroparasites, protozoa, etc.), our results suggest that fungicide applications could cause more harm than good. Indeed, if fungicides and disinfectants do not clear both the frog and environment of *Bd*, our field results suggest that they can eventually elevate *Bd* prevalence. Additionally, many broad spectrum pesticides have widespread non-target effects (Jones et al. 2009, Halstead et al. 2014), but the consequences of fungicide exposure on non-target organisms are not fully understood. Although fungicides show promise for controlling *Bd*, the recently discovered chytrid of salamanders *Batrachochytrium salamandrivorans* (Martel et al. 2013), and perhaps even other emerging fungal diseases, such as those of bats, bees, corals, and snakes (Allender et al. 2011, Cameron et al. 2011, Warnecke et al. 2012), for all of the reasons just provided, we encourage greater research on and caution in using fungicides for managing infectious diseases of wildlife. In particular, research is needed to more concretely identify which fungicides can control fungal pathogens without compromising host pathogen resistance mechanisms.

Although synthetic chemicals provide an enormous value to society, they also have had many unintended consequences (Pimentel et al. 1973). Examples include the paradox of enrichment, where fertilizers reduce crop yields (Rosenzweig 1971) and cases where insecticides cause greater pest outbreaks by reducing natural biocontrol (Desneux et al. 2007, Douglas et al. 2015). Here, we provided yet another example to this growing list. Fungicides paradoxically increased fungal loads and fungal-induced mortality of amphibians. These findings highlight the importance of understanding the role of multiple simultaneous and sequential stressors in biodiversity declines and disease emergences and the need to comprehensively understand the complex effects that chemicals can have on ecosystems to avoid inadvertent and undesirable ramifications.

## Acknowledgements

We thank D. L. Calhoun, S. Paschke, and K. Smalling for feedback on this manuscript. This research was supported by grants from the National Science Foundation (EF-1241889), National Institutes of Health (R01GM109499, R01TW010286), US Department of Agriculture (NRI 2006-01370, 2009-35102-0543), and US Environmental Protection Agency (CAREER 83518801) to J.R.R., and The University of Tampa’s Dana Faculty Development Grant to T.A.M. The field research was supported by the US Geological Survey’s Amphibian Research and Monitoring Initiative. Any use of trade, firm, or product names is for descriptive purposes only and does not imply endorsement by the U.S. Government. W.A.B. did not materially contribute to the model application described in this publication.

## Supplementary Figures

**Fig. S1.**
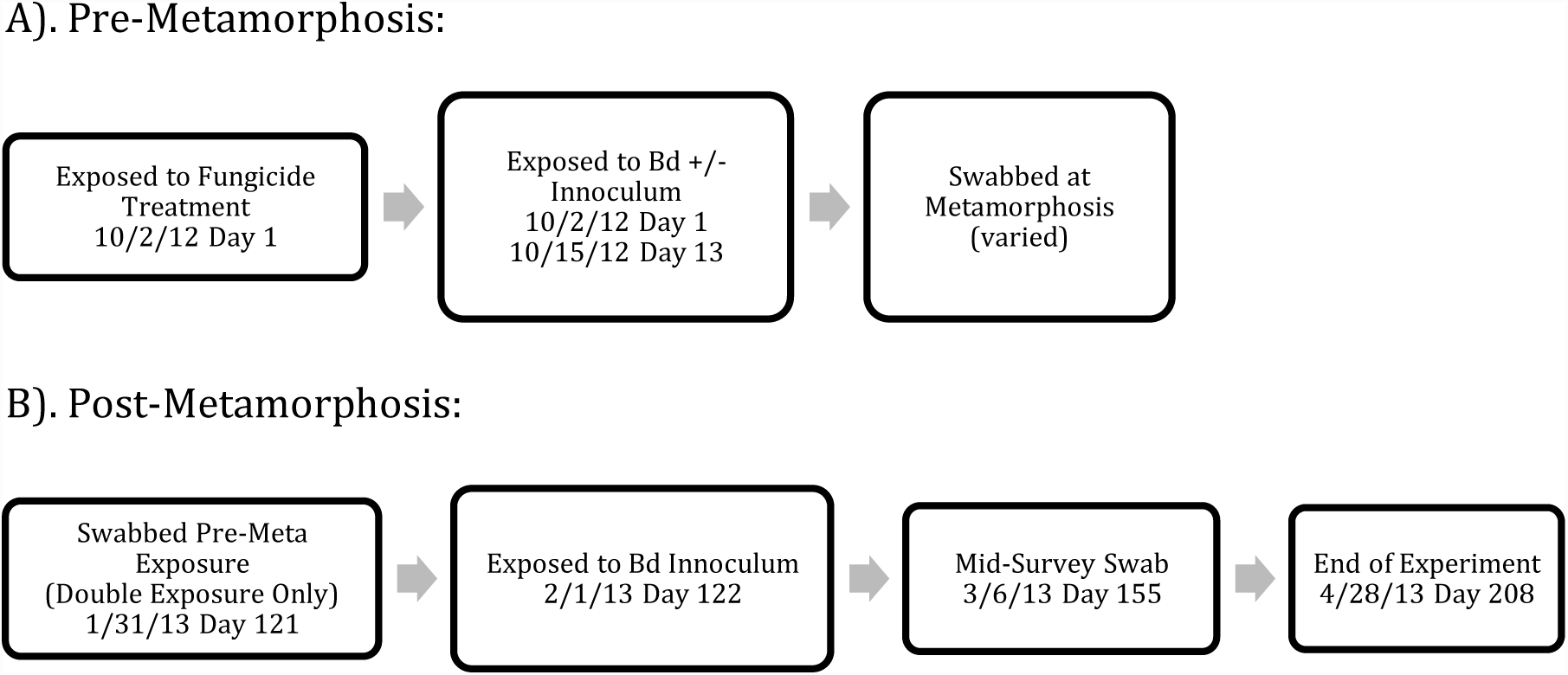
Timeline of events in the lab experiment. **A)** Tadpoles were exposed to a fungicide treatment and Bd treatment on day 1 of the experiment. A follow up *Bd* treatment was administered on day 13 with fungicide treatments reapplied weekly. As animals metamorphosed, they were swabbed. Any tadpoles that did not metamorphose by 12/18/12 were euthanized and swabbed. **B)**All animals exposed to *Bd*+ inoculum as tadpoles were swabbed pre second exposure to assess *Bd* growth. On day 122 of the experiment all juvenile frogs were exposed to *Bd*. Day 155 all metamorphs were swabbed to assess infections. All animals still alive at the end of the experiment were euthanized and swabbed. When an animal died, regardless of life stage, they were swabbed.

**Fig. S2.**
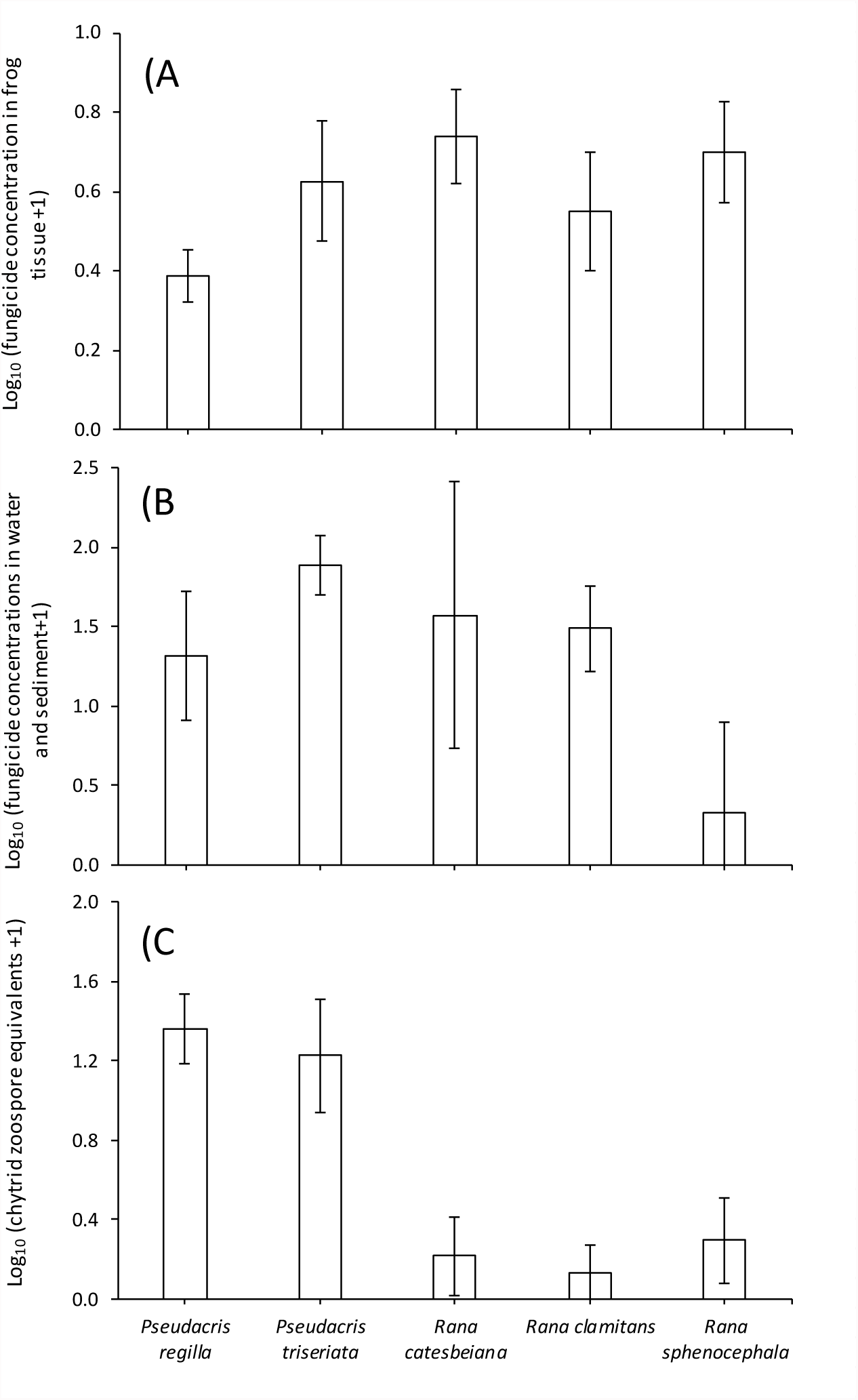
Mean (± 1 SE) fungicide concentrations in frog tissues **(A** and in wetlands *Pseudacris Pseudacris Rana Rana clamitans Rana regilla* and sediment **(B**, and mean chytrid fungal loads **(C** for the five species of frogs *triseriata catesbeiana sphenocephala* (C collected in the field study. Sample sizes for *Pseudacris regilla*, *P. triseriata*, *Rana catesbeiana*, *R. clamitans*, and *R. sphenocephala* are 61, 28, 17, 15, and 17, respectively.

**Table S1.**
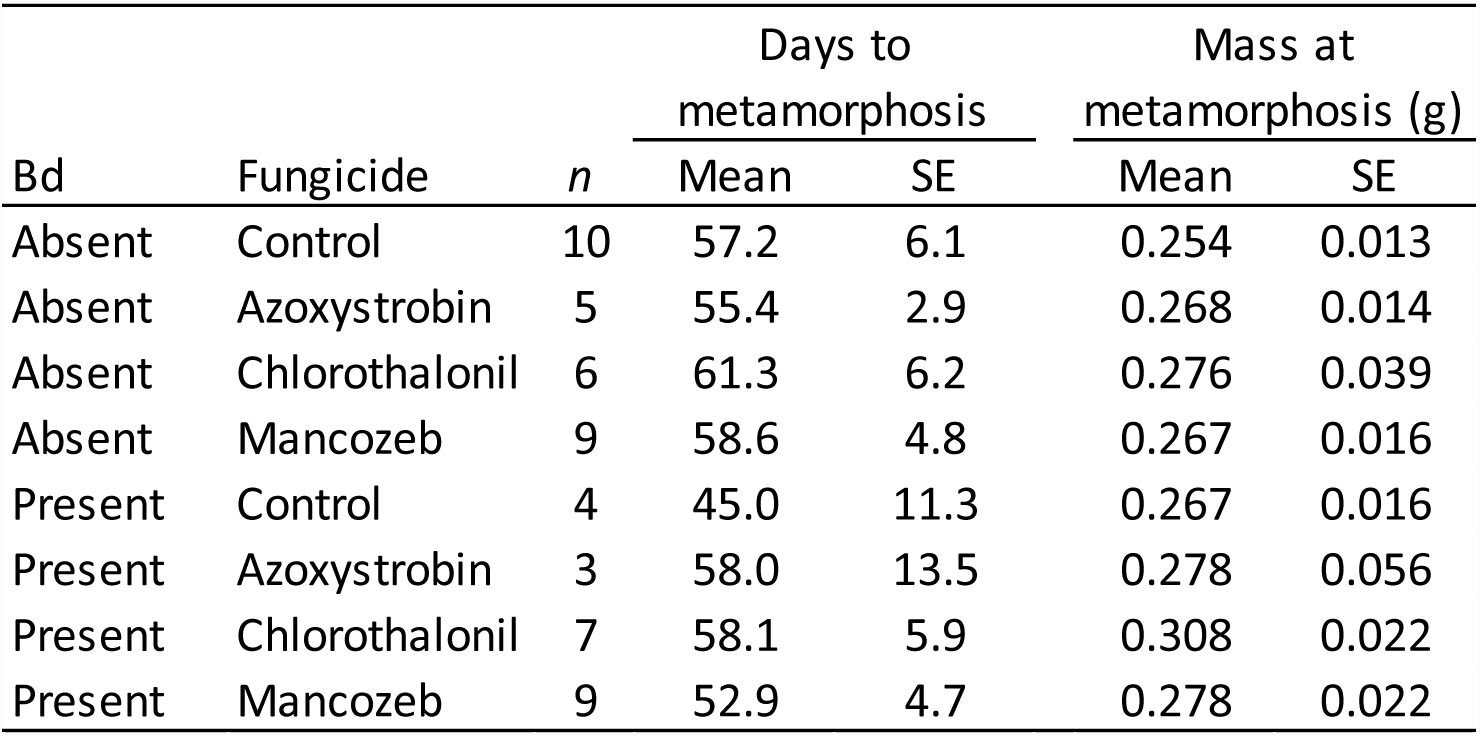
Effects of exposure to fungicides and *Batrachochytrium dendrobatidis* (Bd) on mean time of metamorphosis and size at metamorphosis in Cuban tree frogs.

| Bd | Fungicide | *n* | Days to metamorphosis |  | Mass at metamorphosis (g) |  |
| --- | --- | --- | --- | --- | --- | --- |
|  |  |  | Mean | SE | Mean | SE |
| Absent | Control | 10 | 57.2 | 6.1 | 0.254 | 0.013 |
| Absent | Azoxystrobin | 5 | 55.4 | 2.9 | 0.268 | 0.014 |
| Absent | Chlorothalonil | 6 | 61.3 | 6.2 | 0.276 | 0.039 |
| Absent | Mancozeb | 9 | 58.6 | 4.8 | 0.267 | 0.016 |
| Present | Control | 4 | 45.0 | 11.3 | 0.267 | 0.016 |
| Present | Azoxystrobin | 3 | 58.0 | 13.5 | 0.278 | 0.056 |
| Present | Chlorothalonil | 7 | 58.1 | 5.9 | 0.308 | 0.022 |
| Present | Mancozeb | 9 | 52.9 | 4.7 | 0.278 | 0.022 |

**Table S2.** : Statistical results for the effects of exposure to fungicides and *Batrachochytrium dendrobatidis* (Bd) on time of metamorphosis and size at metamorphosis in Cuban tree frogs.

| Effect | df | Days to metamorphosis |  | Mass at metamorphosis (g) |  |
| --- | --- | --- | --- | --- | --- |
|  |  | *F* | *p* | *F* | *p* |
| Initial mass | 1 | 1.05 | 0.310 | 28.29 | 0.000 |
| Fungicide | 3 | 0.32 | 0.814 | 0.39 | 0.758 |
| Bd | 1 | 1.36 | 0.249 | 0.14 | 0.713 |
| Fungicide * Bd | 3 | 0.29 | 0.834 | 1.03 | 0.388 |

**Table S3.** Effects of fungicide treatments on mean *Batrachochytrium dendrobatidis* (Bd) abundance on Cuban tree frogs when exposed to Bd for the first time as either a tadpole simultaneous with the fungicide exposures or after metamorphosis and after fungicide exposures (i.e. sequential exposures).

| Treatment | Exposed as a tadpole |  |  | Exposed after metamorphosis |  |  |
| --- | --- | --- | --- | --- | --- | --- |
|  | Mean *Bd* |  |  | Mean *Bd* |  |  |
|  | *n* | load | SE | *n* | load | SE |
| Control | 5 | 3.81 | 1.63 | 10 | 1.79 | 0.74 |
| Azoxystrobin | 14 | 4.14 | 0.90 | 10 | 6.12 | 1.20 |
| Chlorothalonil | 10 | 3.24 | 1.15 | 8 | 6.30 | 1.58 |
| Mancozeb | 14 | 1.76 | 0.79 | 14 | 6.22 | 1.08 |

**Table S4.**
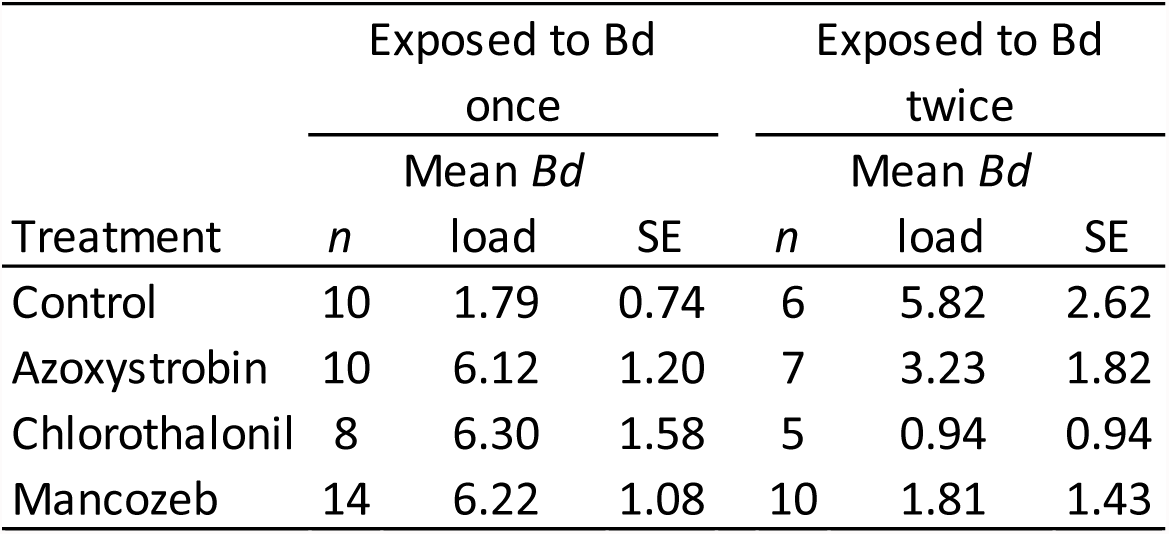
Effects of fungicide treatments on mean *Batrachochytrium dendrobatidis* (Bd) abundance on postmetamorphic Cuban tree frogs exposed to Bd for the first or second time (Le as a tadpole and metamorph).

**Table S5.** Number of wetlands out of 21 sampled where particular fungicides were detected.

| Fungicide | Sediment | Water |
| --- | --- | --- |
| Azoxystrobin | 2 | 4 |
| Boscalid | 3 | 1 |
| Chlorothalonil | 7 | 0 |
| Cyprodinil | 1 | 0 |
| Fludioxinil | 0 | 1 |
| Imazalil | 0 | 1 |
| Fenhexamid | 4 | 0 |
| Propiconazole | 0 | 1 |
| Pyraclostrobin | 11 | 1 |
| Pyrimethanil | 2 | 0 |
| Tebuconazole | 5 | 1 |
| Zoxamide | 2 | 0 |

